# Frequency of change determines effectiveness of microbial response strategies in sulfidic stream microbiomes

**DOI:** 10.1101/2022.04.01.486770

**Authors:** Shengjie Li, Damon Mosier, Xiaoli Dong, Angela Kouris, Guodong Ji, Marc Strous, Muhe Diao

**Author notes:** Corresponding author: Muhe Diao. **Competing Interest Statement** The authors declare no competing interest.

## Abstract

Nature challenges microbes with change at different frequencies and demands an effective response for survival. Here, we used controlled laboratory experiments to investigate the ecological success of different response strategies, such as post-translational modification, transcriptional regulation, and specialized versus adaptable metabolisms. For this, we inoculated replicated chemostats with an enrichment culture obtained from sulfidic stream microbiomes 16 weeks prior. The chemostats were submitted to alternatingly oxic and anoxic conditions at three frequencies, with periods of 1 day, 4 days and 16 days. The microbial response was recorded with 16S rRNA gene amplicon sequencing, shotgun metagenomics, transcriptomics and proteomics. Metagenomics resolved 26 nearly complete genomes of bacterial populations, mainly affiliated with Proteobacteria and Bacteroidetes. Almost all these populations maintained a steady growth rate under both redox conditions at all three frequencies of change. Apparently, oscillating oxic/anoxic conditions selected for generalistic species, rather than species specializing in only a single condition. Rapid (1-day) dynamics yielded more stochasticity, both in community dynamics and gene expression, indicating that bet-hedging might be an effective coping strategy for relatively rapid environmental change. Codon-usage bias, previously associated with copiotrophic and oligotrophic lifestyles, was found to be a powerful predictor of ecological success at different frequencies, with copiotrophs and oligotrophs more successful at a rapid and a slow pace of change, respectively, independent of growth rate.

## Introduction

“The only constant in life is change”, according to the philosopher Heraclitus, ∼500 BC in Ancient Greece. “Changes are in diverse forms, up or down, rigid or flexible, and throughout the whole universe”, as stated in I Ching, an ancient Chinese divination text, ∼1000 BC. Microbes, the smallest and most abundant in numbers among cellular organisms, are coping with change all the time. Microbiomes of the oral and digestive-tract of animals experience dynamics associated with feeding regimes, leading to cycles of feast and famine multiple times per day^1,2^. At even shorter timescales - minutes to seconds - these microbiomes experience periods of oxygen excess or limitation. Cyanobacteria display a progression of gene expression in response to diurnal cycles^3,4^. Seasons dictate change in lakes, with water columns mixing in spring and winter whereas stratifying during summer and fall^5^. At longer timescales, global climate change and eutrophication tip entire ecosystems into different modes of operation^6-8^. Often, change affects redox conditions, which are the topic of this study.

Nature challenges microbes with change of different periodicities, and microbes have developed different coping strategies in response. Post-translational modifications to proteins can (de)activate biochemical pathways quickly, reversibly and with minimal bio-energetic costs^9^. For example, 30% of the yeast proteome is affected by phosphorylation^10^ and a similar extent of the *Escherichia coli* proteome undergoes acetylation^11^. Cyanobacterial rhythms can be governed by phosphorylation of circadian clock proteins^12^. Regulation of gene transcription and translation resets priorities for protein production, remodeling a cell’s proteome in response to a changing situation. Bacterial and archaeal genomes differ from eukaryotic genomes in that subcellular systems are organized in modular gene clusters known as operons. These are expressed under the control of a single response mechanism, such as one/two-component regulators or DNA methylation. In *Escherichia coli*, transcription factors have been shown to regulate operon expression in response to carbon, phosphorus and nitrogen availability^13,14^ Transcriptional control of circadian rhythms is widespread among microbes^15-17^. Interestingly, correlation between mRNA and protein levels is sometimes poor^18,19^. For change at longer timescales, some microbes have developed specialized survival forms, such as spores, that can maintain viability through thousands of years of unfavorable conditions^20^. Evolution is a more general mechanism of adaptation to even slower change, acting over thousands of generations^21^.

Even though it is generally assumed that bacteria are always responsive to their environment, this is not necessarily the case. After all, regulation is associated with trade-offs, such as the bio-energetic costs associated with accelerated turnover of the proteome. Both protein biosynthesis and protein degradation cost energy and consume ATP^22^. Instead of responding to change, microbes may survive a period of unfavorable conditions without adaptation, counting on conditions to become more favorable quickly enough. Alternatively, they may constitutively express a multifunctional proteome that provides answers to different conditions^23^. For example, in bioreactors cycled every 6-12 hours, relatively few proteins were found responsive between oxic and anoxic phases^24,25^. In intertidal sediments, transcription of genes for aerobic respiration and denitrification was unaffected by oxygen concentrations^26^. In tropical forest soils, many taxa displayed sustained activity through rapidly fluctuating redox conditions^27^.

When an ecosystem shifts back and forth between two different redox conditions, will it select for two different, specialized microbiomes, one for each condition? Or will it select for a single, generalist microbiome that functions under both conditions? We hypothesize that this depends on the frequency of change, with generalists successful at a rapid pace and specialists at a slower pace. In generalized microbiomes, will species express a stable proteome that can handle different conditions? Or will they adapt their proteome each time conditions change? We hypothesize that stable expression will occur at a high pace of change, whereas remodeling will happen at a lower pace. Will species have more success with post-translational modification or with transcriptional regulation? Again, we hypothesize this will depend on the pace of change, with a bigger role for transcriptional regulation at a lower rate of change.

To address these hypotheses, we need to pit the different strategies against each other, under defined conditions. For this, we could use an enrichment culture, a consortium of microbes obtained from a single microbiome, assembled in a lab environment relatively recently. Alternatively, we could use a synthetic microbial community, a collection of microbial isolates obtained from culture collections. Here, we opted for an enrichment culture, (a) for avoiding selection effects and evolutionary adaptation associated with long-term lab-cultivation, (b) because more biodiversity in the source community improved the likelihood of representation of varied ecological strategies in the experiment. We used microbiomes sampled from a sulfidic spring as the source community. This community was naturally exposed to redox gradients in space and time and was easily accessible year-round, facilitating future reproduction of the work. No single study will be able to conclusively address the sweeping questions we are asking, so generalization will depend on future studies with a diversity of approaches and source communities.

When a wild microbiome is first transferred to the lab, initial selection may be governed by factors outside the scope of a study’s design - for example, the growth medium may be toxic to some of the microbes present in the natural sample. On the other hand, when enrichment proceeds for too long, evolutionary adaptation to the experimental setup may become a confounding factor^28,29^. To strike a balance between these two, we used a sixteen-week pre-adaptation period in batch-incubations, exposed to alternating redox conditions, followed by the actual experiments conducted in chemostats, for at most 16 generations, at a dilution rate of 0.5 volume changes per day. This corresponds to a doubling time of 1.4 days, much longer than typical for isolated bacteria grown in the lab. We applied three different change regimes (**Fig. 1**). In the first set of triplicated chemostats, cells experienced alternatingly oxic and anoxic conditions about twice per generation. In the second set, redox conditions changed in pace with generation time. In the third set, the cells experienced redox change about once per four generations. We obtained metagenome-assembled genomes (MAGs) for 26 populations and monitored responses by transcriptomics and proteomics. By pitting microbes employing different response strategies against each other, competing for resources in a changing environment, this study set out to falsify our three hypotheses.

**Fig. 1.**
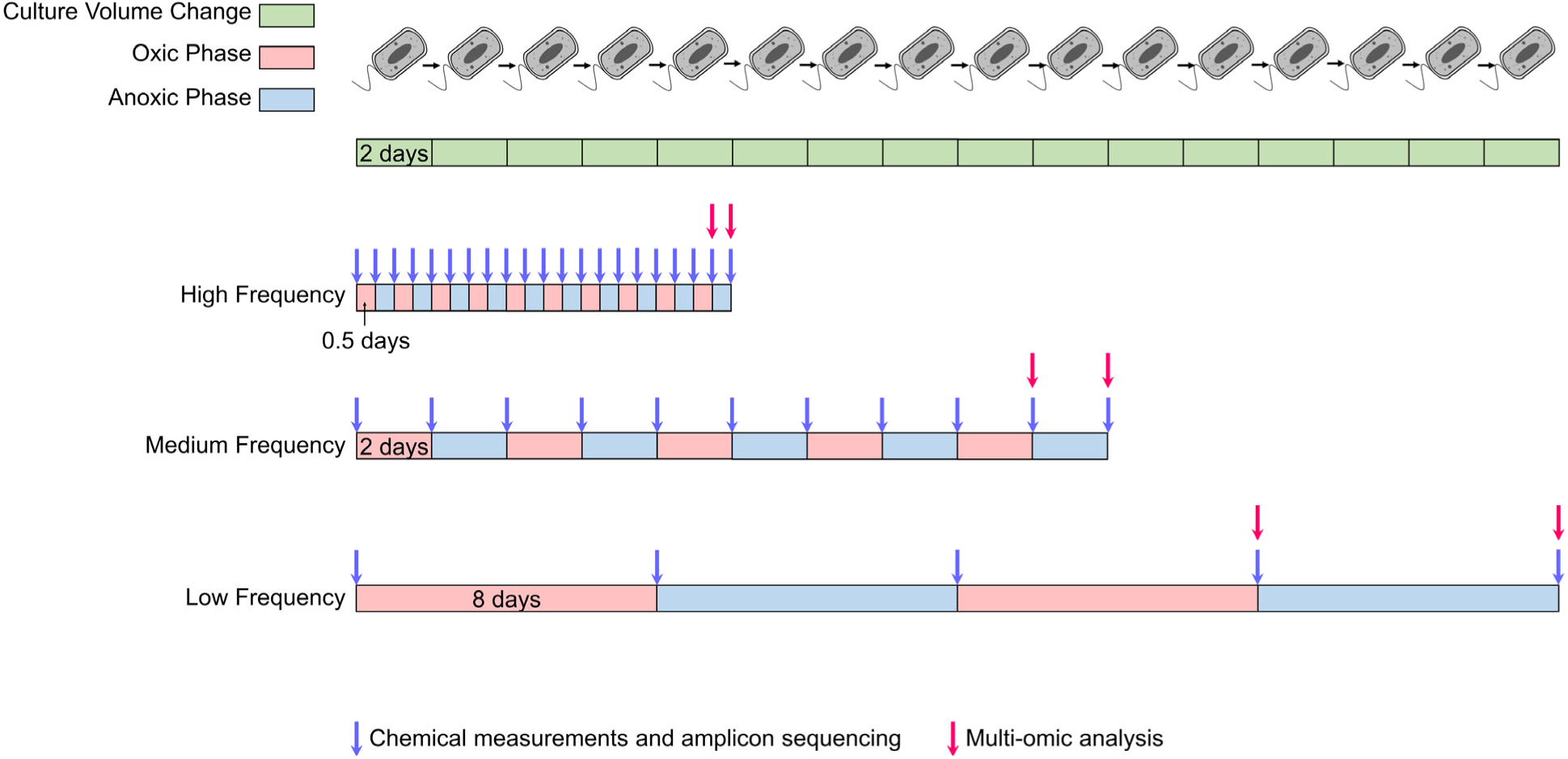
Experimental design of this study. Alternating phases of oxic and anoxic conditions were established in the three sets of triplicated chemostats. The phases differed in length for each set, but the dilution rate was the same for all chemostats.

## Results

Sediment samples were collected from sulfidic streams at Canyon Creek in Canada (Fig. S1a-c). The sediments were naturally exposed to oxygen gradients in space and time (Fig. S1d-j) and both aerobes (e.g. *Thiobacillus, Thiothrix* and *Thiomicrorhabdus*) and anaerobes (e.g. *Geobacter, Desulfocapsa* and *Sulfurovum*) were present in the original sediment community (Fig. S2, Supplementary Table 2). Stream sediments were first pre-adapted to the laboratory in six parallel batch cultures for 16 weeks. Next, we inoculated the combined batch cultures into three sets of chemostat experiments at a dilution rate of 0.5 per day. Each set experienced alternating oxic and anoxic phases, controlled by flushing the chemostats with either air or Argon (Fig. 1). The first set of triplicated chemostats had short phases, so that each generation of microbes experienced oxic/anoxic shifts multiple times. In the second set, each generation experienced only about one shift. The third set had long phases that lasted longer than a single generation. At the end of each oxic and anoxic phase, we determined the outcomes of microbial metabolism based on nutrient concentrations and the community composition with amplicon sequencing. Multi-omics was performed during the final oxic and anoxic phase, to investigate the selected response strategies in more detail (Fig. 1).

### Outcomes of microbial metabolism

Rhythms in nutrient concentrations proceeded in pace with shifts in air and Argon flushing (Fig. 2). This showed that (a) the shifts in redox conditions were successfully established and (b) the enriched bacteria were responding to the shifting conditions. Acetate was always fully consumed during oxic phases. During anoxic phases, it accumulated, up to 5 mM in low-frequency experiments. Sulfate concentrations were stable at high- and medium-frequency. At low-frequency, sulfate accumulated during oxic phases. Sulfide remained undetectable during oxic phases, and accumulated, up to 1.5 mM, during anoxic phases at low-frequency. Nitrate was mostly used up during both oxic and anoxic phases (Supplementary Table 4). These results indicated occurrence of aerobic respiration, (aerobic) denitrification, sulfide oxidation and cysteine metabolism. At low-frequency, total cellular biomass was 3.9±1.5 times higher during oxic phases than anoxic phases (with DNA as a proxy for biomass, Supplementary Table 4). At medium-frequency, oxic biomass was 2.6±1.7 times higher than anoxic biomass. No significant differences between oxic and anoxic biomass were observed at high-frequency. Because aerobic metabolism provides more energy, biomass was expected to be higher at the end of oxic phases than anoxic phases. In addition, cells growing slower than a chemostat’s dilution rate may also be washed out during anoxic periods, reducing biomass. Not all results are easily explained. For example, where ammonia was stable around 0.5 mM at high- and medium-frequency, it accumulated up to 2 mM during oxic phases at low-frequency.

**Fig. 2.**
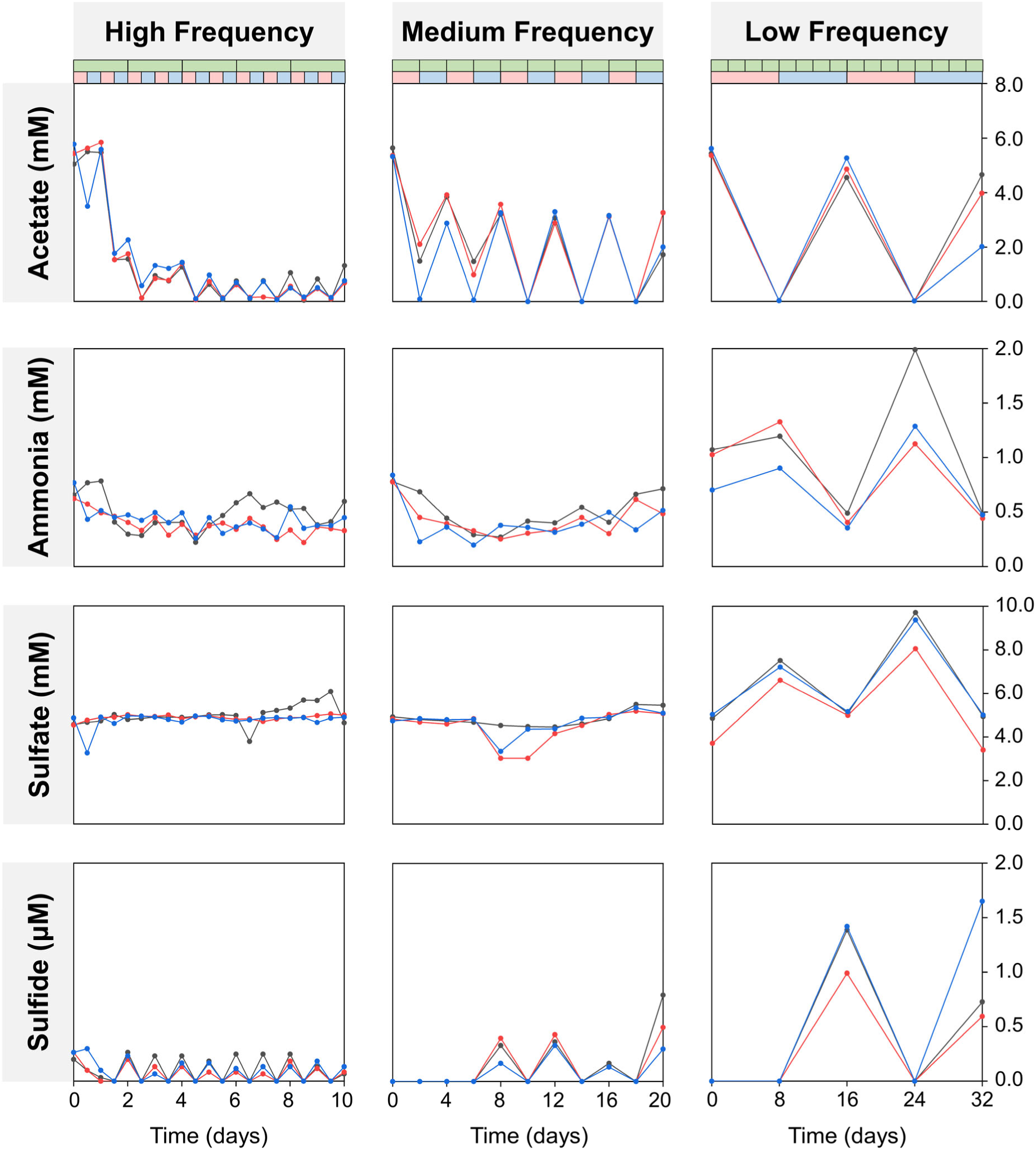
Concentrations of acetate, ammonia, sulfate and sulfide during oxic and anoxic phases at different frequencies. Triplicates are indicated by black, red and blue lines and symbols. The green bar at the top shows chemostat dilutions. In the second bar, oxic and anoxic phases are shown in red and blue, respectively.

### Community dynamics

16S rRNA gene amplicon sequencing showed that by the end of the 16-week pre-adaptation batch cultures, 92 out of 3071 stream populations, each represented by a single amplicon sequence variant (ASV), were still present (Supplementary Table 2). Of 119 abundant stream populations (relative sequence abundance in situ > 0.1%), 34 were still present at the end of the pre-adaptation. Thus, pre-adaptation outcomes were co-determined by in-situ abundance. Although our study did not aim to reproduce sulfidic-stream microbiomes in the lab, we still observed significant representation. Amplicon sequencing also showed that at least 199 populations in total were present at the end of the pre-adaptation/chemostat inoculation. This biodiversity was the starting point for subsequent selection. Selection in chemostats was very effective, as within two days, a simple microbial community established itself in each experiment. These communities featured the same ten abundant populations (ASVs), making up > 88% of relative sequence abundance across all chemostat experiments and replicates (Fig. 3a, Supplementary Table 2). Three of these ten populations, **3** *Thiobacillus*, **9** *Arenimonas* and **10** *Brevundimonas* were ubiquitous in stream microbiomes. Among the stream’s ten most abundant populations, three were represented in the chemostats, including **3** *Thiobacillus* (Fig. S2). Among these ten most abundant populations, success differed between treatments.

**Fig. 3.**
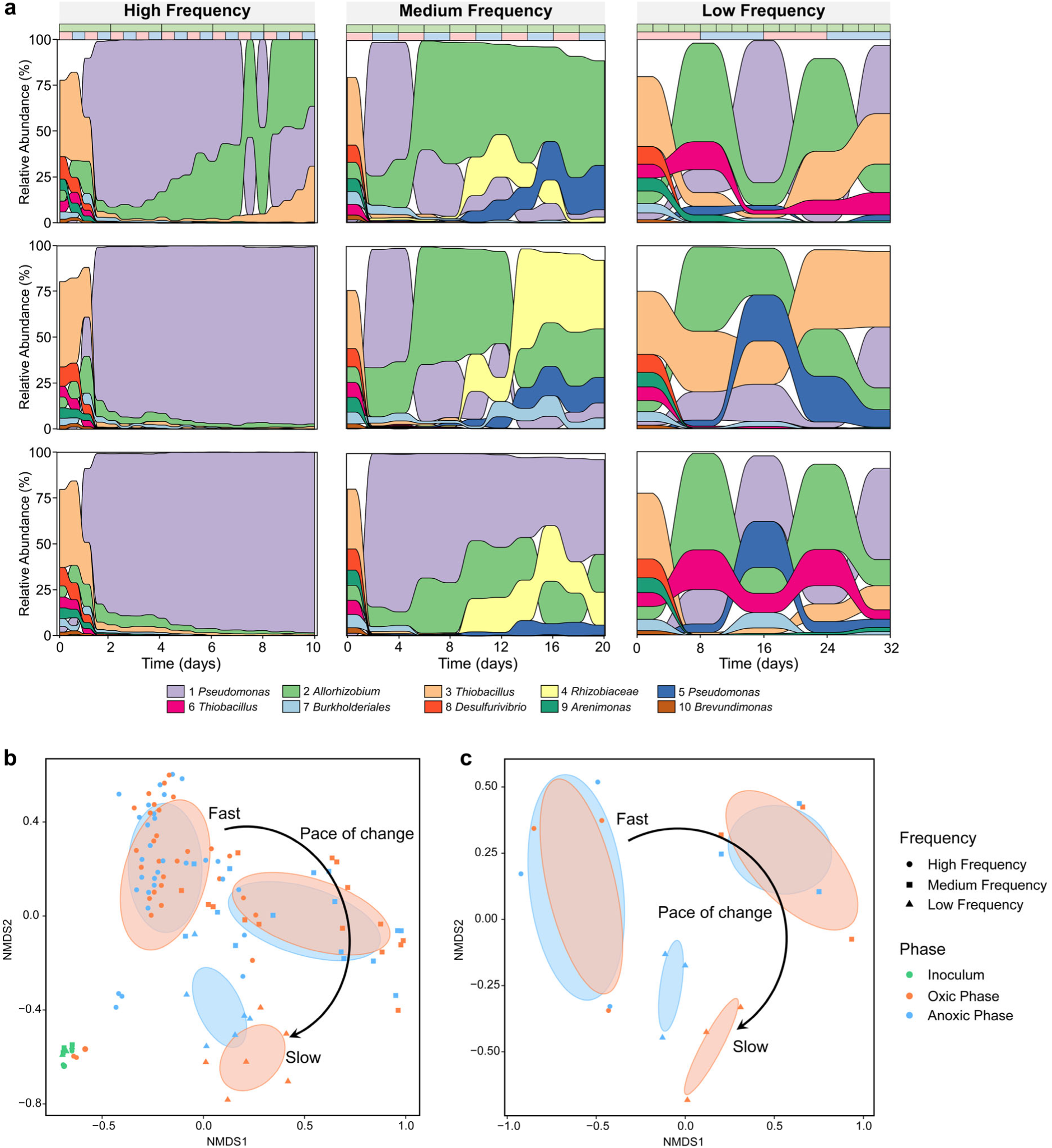
Community dynamics in the chemostats. **a**, Change in relative sequence abundances of the ten most abundant populations (amplicon sequence variants, ASVs) in chemostat incubations, based on 16S rRNA gene amplicon sequencing. Outcomes of triplicated experiments are shown individually for each frequency. The green bar at the top enumerates chemostat dilutions. In the second bar, oxic and anoxic phases are shown in red and blue respectively. **b**, Non-metric multidimensional scaling (NMDS) (based on Bray-Curtis distances) of all samples collected along the chemostat incubations and **c**, only samples collected during the final oxic and the final anoxic phases. Colored ellipses show variation among samples of the two phases of each treatment obtained with the ‘ordiellipse’ function from the ‘vegan’ package in R.

At high-frequency, the community composition was relatively stable between oxic and anoxic phases, with **1** *Pseudomonas* dominating communities in two of the replicates. At low-frequency, community compositions oscillated in tune with oxic and anoxic conditions. For example, **1** *Pseudomonas* and **2** *Allorhizobium* were more abundant during anoxic and oxic phases respectively. This was the only treatment where community differences between phases were larger than between replicates, as shown by nonmetric multidimensional scaling (Fig. 3b,c).

Differences between replicates were large at high and medium-frequency and small at low-frequency. Frequency of change was more important in shaping community structure than the occurrence of oxic or anoxic conditions. For example, **4** *Rhizobiaceae* was most successful at medium-frequency, whereas **3** *Thiobacillus* was mainly observed at low-frequency. All abundant populations appeared to be able to cope with both redox conditions, indicating they might be capable of both aerobic and anaerobic metabolism.

### Physiology and growth of enriched populations

To investigate the metabolic potential and lifestyle of the enriched populations more closely, shotgun metagenomes were sequenced for samples collected at the end of the final oxic and anoxic phases of each chemostat experiment. The metagenomes were assembled and binned into MAGs. We obtained 26 MAGs in total, including 18 MAGs with completeness > 90% and contamination < 10% (Supplementary Table 5). The total abundance of the 26 MAGs accounted for over 99% of sequenced DNA in all samples (Fig. 4a, Supplementary Table 5). Although the metagenomes were obtained at the end of the experiments, the MAGs were associated with populations active throughout chemostat selection. Only populations that were completely unsuccessful and disappeared from the experiment entirely, such as ASV **8** *Desulfurivibrio*, were not represented among MAGs. Community composition based on 16S and shotgun sequencing were consistent, but relationships between ASVs and MAGs were not always one to one. For example, ASV “**1** *Pseudomonas*” was associated with two MAGs, “Pseudomonas A” and “Pseudomonas C”.

**Fig. 4.**
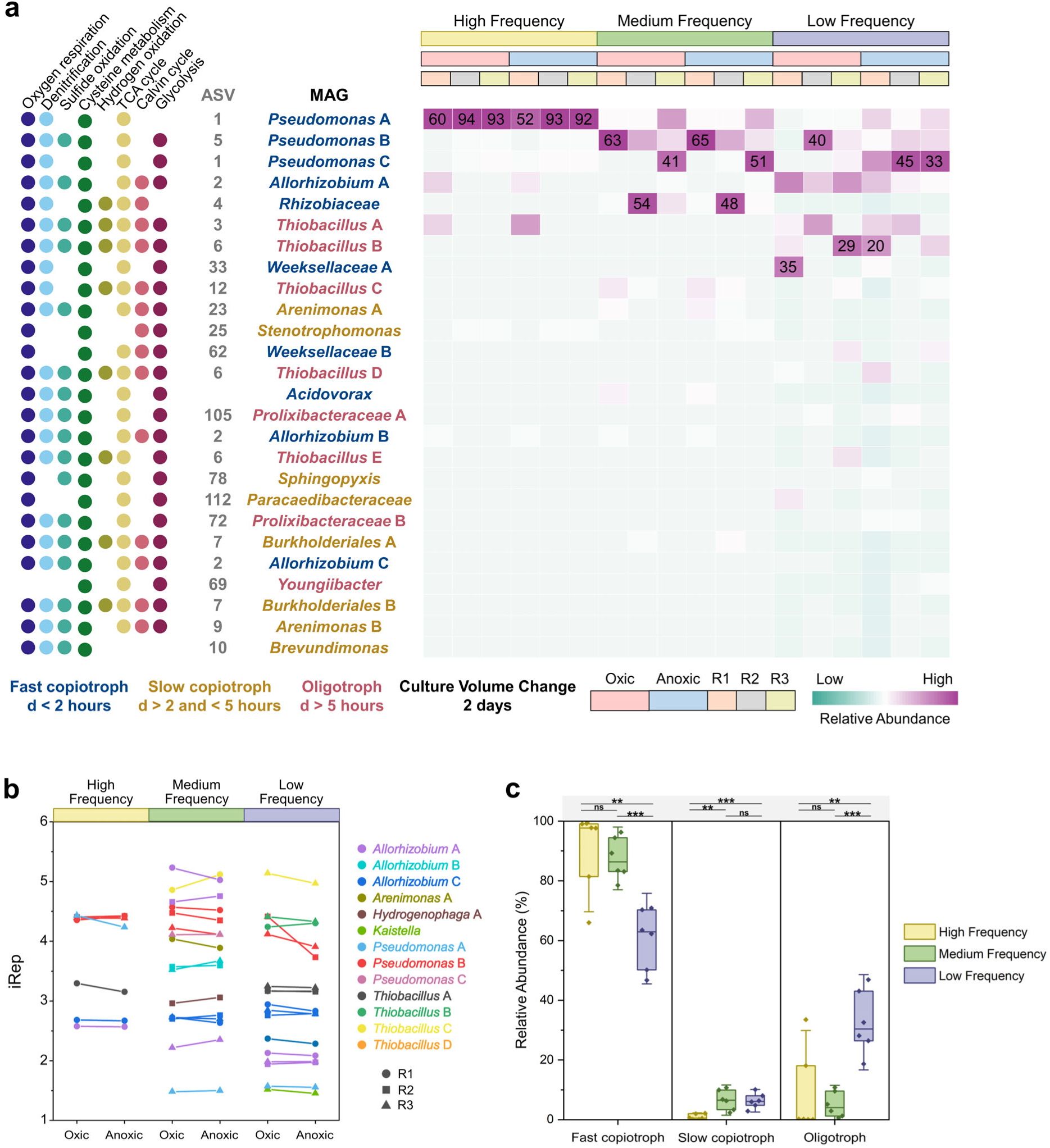
Enriched populations associated with metagenome-assembled genomes (MAGs) in the three sets of chemostats during the final oxic phase and the final anoxic phase. **a**, Metabolic potential, taxonomy and relative sequence abundance of the populations (Supplementary Tables 5-31). R1, R2 and R3 represent the chemostat triplicates. Fast copiotrophs, slow copiotrophs and oligotrophs are indicated by blue, yellow and pink taxon names, respectively. “d” represents the minimum doubling time predicted by gRodon^33^ (Supplementary Table 33). **b**, iRep values (Supplementary Table 32) of the populations. **c**, Relative abundance of fast copiotrophs, slow copiotrophs and oligotrophs in the three sets of chemostats. Horizontal lines show significant differences determined in t-tests, with P-values <0.01 indicated with “**” and < 0.001 indicated with “***”.

Analysis of gene content of MAGs indicated that 22 out of 26 associated populations selected during chemostat trials were capable of both aerobic and anaerobic growth (Fig. 4a, Supplementary Tables 6-31). Most deciphered metabolic pathways were encoded in > 50% of MAGs, indicating vast functional redundancy among community members. As cysteine was a major source of energy, carbon and sulfur in the medium, it was not surprising that all selected populations encoded cysteine desulfurase in their genomes. Aerobic respiration, denitrification and sulfide oxidation were common and consistent with observed nutrient dynamics (Fig. 2). One metabolic pathway we did not expect to see was the Calvin cycle for carbon fixation: ten genomes contained both ribulose-1,5-bisphosphate carboxylase/oxygenase (RuBisCO) and phosphoribulokinase (PRK)^30^. Apparently, almost half of the enriched populations potentially used carbon dioxide as a carbon source even though organic substrates such as acetate were often present in excess (Fig. 2).

The capacity of most populations to grow both aerobically and anaerobically was supported by stable relative sequence abundance of associated MAGs during oxic and anoxic phases at high- and medium-frequency, in agreement with amplicon sequencing results. However, at low frequency, **1** *Pseudomonas* AC and **2** *Allorhizobium* A were more abundant during anoxic and oxic phases respectively. To compare aerobic and anaerobic growth rates of individual populations, we calculated the peak-to-trough (PTR) ratio of sequencing depth for each MAG during oxic and anoxic conditions with iRep^31^. A high growth rate is associated with a high genome replication rate, resulting in a high PTR ratio. Although PTR is a poor proxy for growth rate when comparing different species^32^, it works well for comparing growth rates of the same species across different samples^31^. We found PTR ratios did not differ significantly between phases, even for **1** *Pseudomonas* AC and **2** *Allorhizobium* A at low frequency (Fig. 4b, Supplementary Table 32). This indicated that aerobic and anaerobic growth rates were similar. Overall, analysis of PTR ratios supported the conclusion that dynamic conditions selected for species that coped well with both oxic and anoxic conditions. Even in the case of **1** *Pseudomonas* AC and **2** *Allorhizobium* A at low-frequency, the observed changes in abundance could be explained with only minor differences in growth rate.

In contrast to redox state, frequency of change did select for specific populations, in agreement with amplicon sequencing results. For example, **1** *Pseudomonas* AC, **4** *Allorhizobium* B and **3**,**6** *Thiobacillus* AB were most successful at high-, medium- and low-frequency respectively. To explain this outcome, we investigated the codon usage bias of the MAGs with gRodon^33^ (Supplementary Table 33). Strong codon usage is associated with rapid growth, but could also facilitate rapid gene expression in response to environmental cues. According to the predicted minimum doubling times by gRodon, we grouped the populations to fast copiotrophs (< 2 hours), slow copiotrophs (> 2 and < 5 hours) and oligotrophs (> 5 hours) (Fig. 4c, Supplementary Table 33). The chemostats selected for a mix of copiotrophs and oligotrophs (Fig. 4c). However, fast copiotrophs were more successful at high- and medium-frequency than low-frequency, while slow copiotrophs were more successful at medium- and low-frequency, and oligotrophs were more successful at low-frequency (Fig. 4c). Thus, codon usage bias was identified as an important predictor of success in coping with change of different frequencies.

### Transcriptional and translational regulation

Metatranscriptomics and proteomics were used to determine changes to each population’s gene expression from the final oxic phase to the final anoxic phase (Fig. 5a, Supplementary Tables 34-41). Genes involved in all investigated metabolic categories, including the Calvin cycle, were active during both oxic and anoxic phases. At all three frequencies, genes for aerobic respiration were more actively transcribed during oxic phases, while denitrification genes were more active during anoxic phases. For some subsystems responses depended on frequency. For example, hydrogen oxidation by NiFe hydrogenases was more active during the oxic phase at low-frequency and during the anoxic phase at high-frequency. Most subsystems showed no significant differences in proteomes.

**Fig. 5.**
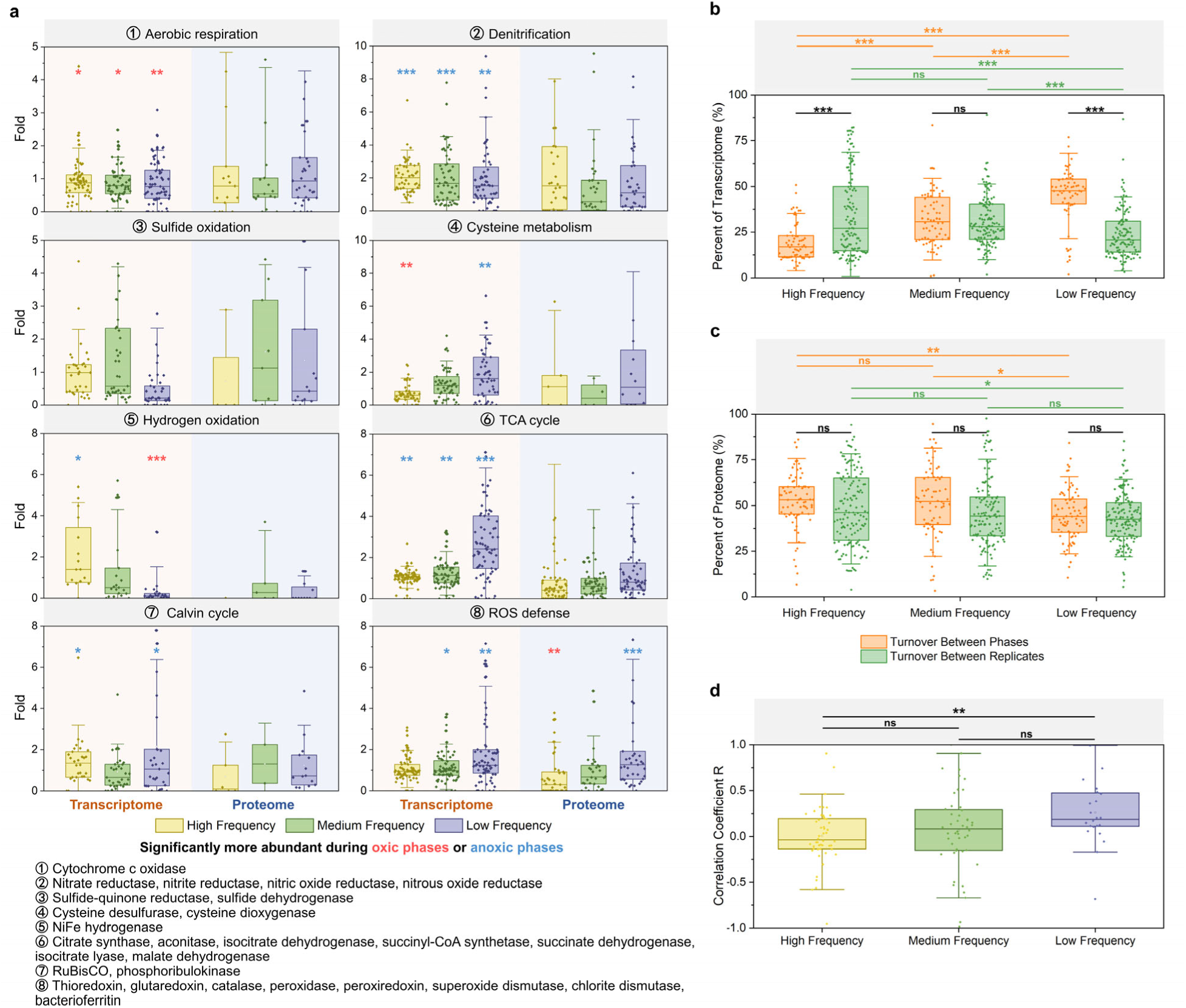
Change in transcriptomes and proteomes in the three sets of chemostats. **a**, Fold change in activity of genes associated with key metabolic subsystems (Supplementary Table 34-41) from the final oxic phase to the final anoxic phase. The circled numbers indicate the corresponding enzymes involved in the metabolic subsystems. Each data point is associated with one of 26 MAGs. Results are shown for transcriptomes (left) and proteomes (right), each at the three different frequencies of change. Significances, determined with t-tests, are indicated, with P-values <0.05 as “*”, <0.01 as “**” and < 0.001 as “***”. **b**, Turnover of transcriptomes across phases and replicates (Supplementary Table 42). Each dot shows overall transcriptome turnover between phases and replicates for a single MAG. **c**, Turnover of proteomes across phases and replicates (Supplementary Table 42). Each dot shows overall proteome turnover between phases and replicates for a single MAG. **d**, Pearson correlation coefficients of transcriptome differences and proteome differences between phases for a single MAG (Supplementary Table 43). Each dot shows the correlation between the transcriptome and the proteome for a single MAG. Horizontal lines in **b, c** and **d** show significant differences determined with t-tests.

To explore the role of gene expression in adaptation more broadly than these categories, we calculated the “turnover” of each population’s entire transcriptome and proteome across phases and frequencies. The turnover is the percentage of the transcriptome/proteome that differs between two samples. To get a sense of the experimental noise and natural variability, we compared the turnover from the oxic to the anoxic phase of individual replicates to the turnover between two replicates at the same phase (oxic or anoxic).

Whereas transcriptome “turnover” between replicates increased with increasing frequency, transcriptome turnover between phases decreased with increasing frequency (Fig. 5b, Supplementary Table 42). Apparently, exposure to high-frequency change led to higher natural variability in transcriptomes. Only at low frequency was transcriptome turnover associated with regulation (47.9±15.4%, n=63) higher than stochastic differences between replicates (24.0±13.4%, n=118). Transcriptome turnover was lower than cell turnover at all three frequencies, indicating that organisms may not have actively degraded old transcripts (messenger RNA), but only responded to change by adding new ones.

Surprisingly, proteomes showed a different trend (Fig. 5c, Supplementary Table 42): Here, turnover between replicates and phases both decreased with decreasing frequency of change. For all three frequencies, there were no significant differences between the impact of natural variability and regulation. Proteome turnover between phases was similar to transcriptome turnover between phases at low-frequency, but much higher than transcriptome turnover between phases at high-and medium-frequency. Proteome turnover was lower than cell turnover at medium- and low-frequency, but larger than cell turnover at high-frequency. This indicated active proteome remodeling (degradation of old proteins). Alternatively, and perhaps more likely, these results could also be explained by active use of post-translational modifications (PTMs, see below) at high frequency. Because many PTMs are unknown and not in our databases or might have been lost during sample processing, proteins with PTMs may not have been identified, artifactually increasing protein turnover numbers.

The coherence of transcriptional and translational responses was determined by calculating the Pearson correlation coefficient of mRNA and protein differences between phases (Fig. 5d, Supplementary Table 43). Significantly higher correlations were observed at low-frequency (0.26±0.36, n=24) than high-frequency (0.0053±0.31, n=50) and medium-frequency (0.066±0.43, n=48). Thus, the consistency of gene expression between transcription and translation increased as frequency decreased. Coherence between the transcriptome and proteome was only observed when the pace of change was lower than the generation time.

### Post-translational modifications

Post-translational modification (PTM) is a mechanism for rapidly activating or suppressing a protein’s function. Phosphorylation and acetylation are two commonly observed PTMs^9^. In total, we observed 2,320 phosphorylation events and 2,003 acetylation events with high confidence across all 18 replicates and conditions (Fig. S3, Supplementary Tables 44,45). We observed more phosphorylated proteins at medium-frequency (0.38∼1.9% of detected proteins) compared to the other two frequencies (0.10∼0.77% of detected proteins). Phosphorylation was mainly observed for enzymes involved in central metabolism (the TCA cycle and glycolysis), such as Enolase, Malate dehydrogenase and Phosphoglycerate kinase. Acetylated proteins were detected in similar amounts at the three frequencies (0.17∼0.96% of detected proteins) and were more often observed for proteins associated with the cell envelope, including membrane proteins, flagellar biogenesis, and regulators, such as molecular sensors and two-component response regulators.

## Discussion

We hypothesized that at high-frequency, redox change would select for a single, generalist microbiome capable of coping with both oxic and anoxic conditions. In contrast, at low-frequency we expected selection of two specialized microbiomes. This hypothesis could not be rejected, because the amplicon data clearly showed diverging aerobic and anaerobic microbiomes at low-frequency. At the same time, even at low-frequency, populations displayed relatively stable growth independent of redox conditions, as shown by co-expression of genes for both aerobic and anaerobic metabolism, stable peak-to-trough ratios in genome sequencing depth and persistent relative sequence abundances, with few exceptions at low-frequency. Note that our low-frequency experiment featured eight-day long oxic and anoxic phases, much longer than common natural oscillations such as day/night cycles and feeding regimes. Thus, even though specialization of microbiomes was detectable, the enriched microbiomes were overwhelmingly generalist. At an even slower pace of change, selection of specialized microbiomes will eventually proceed, as reported in previous studies addressing, for example, seasonal change^34,35^.

We also hypothesized that at high-frequency, gene expression would be stable throughout oxic and anoxic conditions. In contrast, at low-frequency we expected changes in gene expression. This hypothesis was falsified. For example, expression of genes involved in denitrification was higher during anoxic conditions, independent of frequency. In addition, overall, gene expression was more variable at high-frequency. This variability was not associated with changing conditions but was stochastic, caused by differences between replicates. High-frequency change was also associated with higher variability in community composition between replicates, as shown by amplicon sequencing. Apparently, high-frequency change increased variability, both at the level of populations and their phenotypes. This variability/stochasticity might result from adaptation in the form of “bet-hedging”^36,37^. In contrast, at low-frequency, variability in gene transcription could be mainly explained by redox state. This was also the only frequency at which transcriptomes and proteomes were overall consistent with each other, showing effectiveness of transcriptional regulation. Thus, although frequency of change did influence the effectiveness of transcriptional regulation as hypothesized, outcomes differed between metabolic subsystems and stochasticity was found to be important, especially at high-frequency.

Our third hypothesis was that post-translational modification (PTM) and transcriptional regulation would be more important at high- and low-frequency, respectively. This hypothesis could not be rejected. As already discussed, transcriptional regulation was indeed found to be more pronounced at low-frequency. We did not obtain direct evidence for increased importance of PTM at high-frequency. This might be explained by losses of PTMs during sample processing in combination with occurrence of a large untested diversity in potential PTMs. However, indirect evidence in the form of very high proteome turnover at high-frequency experiments hinted at a role of PTM in coping with rapid change. Occurrence of unknown PTMs would lead to a failure to correctly identify peptides during proteomics and could explain large differences in proteomes between samples. Dedicated approaches to quantify PTMs will be needed to address this hypothesis more conclusively^38,39^.

In addition to addressing the hypotheses, the experiments also yielded unforeseen outcomes. First, we observed that codon usage bias, previously associated with oligotrophic and copiotrophic lifestyles, predisposes oligotrophs and copiotrophs to cope well with slow and fast change, respectively. In our experiments, success was not determined by differences in growth rate, as the growth rate was the same across all experiments. Because strong codon bias could enable copiotrophs to more rapidly respond to change, this outcome can still be easily understood. Second, expression of genes involved in carbon dioxide fixation might have contributed to the success of many of the enriched species, even though organic carbon in the form of acetate was often present in excess.

To what extent can results of this study be generalized? At least, some key premises of the research were validated experimentally. The source microbiome contained a mixture of aerobes and anaerobes, naturally exposed to gradients of redox conditions in space and time. After pre-adaptation in the lab, the community still featured at least 199 amplicon sequence variants that were available for chemostat selection to act upon. That is a much greater starting diversity than would be feasible using a synthetic community approach. Selection during chemostat experiments was very effective as anticipated. The outcomes of selection differed between treatments, and could be explained by a fundamental property, codon usage bias, which controls the speed of the transcriptional response. Nutrient concentrations clearly showed the outcomes of aerobic and anaerobic metabolism during oxic and anoxic phases, respectively. Thus, the deployed ecological strategies worked and showed that the enriched microbial communities successfully coped with change during the experiments. The enriched communities were mainly made up of Proteobacteria, Bacteroidetes and Firmicutes, globally distributed, ecologically successful bacterial phyla. These bacteria were only in the lab for sixteen weeks, a limited time to evolve compared to isolated strains, and recorded phenotypes were more likely to be reflective of the natural situation. Thus, the merits of taking an enrichment approach to address the question still appear strong in retrospect. In conclusion, we incubated a sulfidic stream microbiome in replicated chemostats subjected to oxic/anoxic change at different frequencies. We found that generalists capable of both oxic and anoxic metabolism were more successful than specialists and that these bacteria co-expressed genes for aerobic and anaerobic metabolism continuously, independent of redox state. High and low frequency of change was found to select for copiotrophs and oligotrophs respectively and this defines a novel aspect of these ecological niches. Future studies with different approaches and source microbiomes will show if these findings can be generalized.

## Materials and methods

### Sampling, pre-incubation and chemostat incubation

Sediment samples were collected at six sampling sites from sulfidic streams at Canyon Creek, Canada (50.95159°N, 114.55951°W) on March 5th, 2020. 60 g of the mixed sediments were inoculated into six 1 L serum bottles with 600 mL fresh medium. The medium contained MgCl_2_ • 6H_2_O (2 mM), KH_2_PO_4_ (0.7 mM), CaCl_2_ (0.9 mM), NH_4_Cl (1.9 mM), Na_2_SO_4_ (2.5 mM), NaNO_3_ (1 mM), sodium acetate (2 mM), FeCl_2_ (10 mM), NaHCO_3_ (20 mM), trace element solution (1 mL/L) and vitamin solution (1 mL/L). Trace element solution contained (per liter) titriplex III (EDTA) (0.5 g), FeSO_4_ • 7H_2_O (0.2 g), ZnSO_4_ • 7H_2_O (0.01 g), MnCl_2_ • 4H_2_O (0.003 g), H_3_BO_3_ (0.03 g), CoCl_2_ • 6H_2_O (0.02 g), CuCl_2_ • 2H_2_O (0.001 g), NiCl_2_ • 6H_2_O (0.002 g) and Na_2_MoO_4_ • 2H_2_O (0.003 g). Vitamin solution contained (per liter) biotin (0.1 g), 4-aminobenzoic acid (0.5 g), calcium pantothenate (0.1 g), thiamin (0.2 g), nicotinic acid (1.0 g), pyridoxamine (2.5 g) and vitamin B12 (0.1 g). The six serum bottles were incubated in the dark, at room temperature in a shaker for 16 weeks, for lab acclimatization. During acclimatization, the bottles were alternately incubated with and without oxygen for one week (8 oxic phases, 8 anoxic phases). At the beginning of each oxic phase, the bottles were flushed with helium. Then, about 7% oxygen (final concentration) was injected into each bottle with a syringe. At the beginning of each anoxic phase, the bottles were only flushed with helium. Sodium acetate (2 mM), NaNO_3_ (1 mM) and NaHCO_3_ (20 mM) (final concentrations) were added to the bottles every 4 weeks.

After 16 weeks, the acclimatized cultures were used to inoculate chemostat incubations. For this, the six cultures were first mixed together and then used as inoculum for three sets of triplicated chemostats. 100 mL of mixed acclimatized culture were added to each 1 L chemostat with 900 mL fresh medium. The fresh medium contained MgCl_2_ • 6H_2_O (2 mM), KH_2_PO_4_ (0.7 mM), CaCl_2_ (0.9 mM), NH_4_Cl (1.9 mM), Na_2_SO_4_ (5 mM), NaNO_3_ (2 mM), sodium acetate (5 mM), NaHCO_3_ (20 mM), trace element solution (1 mL/L) and vitamin solution (1 mL/L) and L-cysteine (5 mM). L-cysteine solution was filter sterilized in an anaerobic chamber and kept anoxic before it was added to the medium bottles. The final pH was 6.5 - 7.5.

Each chemostat setup consisted of a 1 L medium (feed) bottle, a 1 L magnetically stirred culture bottle and an effluent collection bottle (Fig. S4). Fresh medium was pumped from the medium bottle to the culture bottle at a rate of 0.5 L per day (one volume change per 2 days). The total culture volume of the culture bottles was maintained at 1 L by pumping out the excess culture volume to the effluent collection bottle. All culture bottles of the chemostats were covered with aluminum foil and stirred at 300 rounds per minute with a magnetic stir bar. Just like in the pre-incubation, the chemostats experienced alternatingly oxic and anoxic conditions. During oxic phases, 10 mL/min air was supplied to the medium bottle and the culture bottle. During anoxic phases, 10 mL/min Argon was supplied to the medium bottle and the culture bottle. 2 mM FeCl_2_ was added directly to the culture bottles at the beginning of every oxic phase. Chemostats were started with an oxic phase and ended with an anoxic phase.

There were 3 treatments for the chemostat incubations, high-frequency, medium-frequency and low-frequency (Fig. 1). For each treatment, a set of triplicated chemostats was run. The nine chemostats were operated independently in parallel with the same inoculum. For high-frequency experiments, each phase lasted for 0.5 days and the total culture time was 10 days (10 oxic phases and 10 anoxic phases, 5 culture volume changes). For medium-frequency experiments, each phase lasted for 2 days and the total culture time was 20 days (5 oxic phases and 5 anoxic phases, 10 culture volume changes). For low-frequency experiments, each phase lasted for 8 days and the total culture time was 32 days (2 oxic phases and 2 anoxic phases, 16 culture volume changes).

Culture samples were collected immediately at the beginning of the incubations and at the end of every phase. The samples were centrifuged at 5,000 rpm for 10 min. 0.2μm-filtered supernatants were used for chemical measurements and cell pellets were used for DNA, RNA and protein extractions. The workflow of the experiment and data analysis were illustrated in Fig. S5.

### Chemical measurements

Sulfide, ammonia, ferrous iron and nitrite were determined with an Evolution 260 Bio UV-Visible Spectrophotometer (Thermo Scientific, CA, USA). Ammonia was measured by the indophenol reaction^40^. Nitrite was measured by a reaction with sulfanilamide and N-(1-naphthyl)ethylenediamine^41^. Sulfide was fixed with zinc acetate and determined by a reaction with dimethylparafenyldiamine and Fe(NH4)(SO4)2 • 12 H2O as previously described^42^. Ferrous iron was measured with the ferrozine method^43^. Acetate was quantified by high-performance liquid chromatography (HPLC) using a Thermo RS3000 HPLC fitted with a Kinetex 2.6 μm EVO C18 100 Å LC column, a Thermo RS3000 pump and an UltiMate 3000 fluorescence detector (Thermo Scientific, CA, USA). Nitrate and sulfate were measured by a Dionex ICS-5000 Ion Chromatography System (Thermo Scientific, CA, USA) equipped with an anion-exchange column (Dionex IonPac AS22; 4×250 mm; Thermo Scientific), an EGC-500 K_2_CO3 eluent generator cartridge and a conductivity detector.

### DNA extraction and amplicon sequencing

DNA was extracted from sediments, pre-incubated culture pellets and chemostat culture pellets with the FastDNA SPIN Kit for Soil (MP Biomedicals, Solon, OH, USA). Qubit 2.0 Fluorometer (Invitrogen, CA) was used to estimate the DNA concentration. Amplicon sequencing was performed with the primers A519F (5’-CAGCMGCCGCGGTAA-3’) and Pro805R (5’-GACTACNVGGGTATCTAATCC-3’), targeting both archaea and bacteria^44,45^. PCR systems were prepared with template DNA, the forward and the reverse primers and 2x KAPA HiFi Hot Start Ready Mix (Roche, CA). PCR was performed with the following protocol: an initial denaturation cycle (95°C for 3 min), 25 cycles of denaturation (95°C for 30 s), annealing (55°C for 45 s) and extension (72°C for 60 s), and a final extension cycle (72 °C for 5 min). Triplicated PCR reactions were conducted for each DNA sample and the PCR products were verified by 1% agarose gel electrophoresis. The amplicons were pooled, purified and sequenced with an Illumina Miseq System (Illumina, San Diego, CA) using the 2 × 300 bp MiSeq Reagent Kit v3. Raw data was processed with amplicon sequencing variant (ASV) analysis in MetaAmp^46^. Non-metric multidimensional scaling (NMDS) analysis was performed with the ‘vegan’ package in R v4.0.3. Different groups were labelled with the ‘ordiellipse’ function, which invisibly returns an object that has a summary method that returns the coordinates of centroids and areas of ellipses. A total of 123 samples collected from the sediments and cultures were sequenced, yielding 4,632,919 reads after quality control (4,858 to 169,903 reads per sample).

### Metagenomic sequencing and data analysis

18 samples were selected for metagenomic, metatranscriptomic and metaproteomic analysis (Fig. S6). These samples were collected at the final oxic and anoxic phases of the triplicated treatments (2×3×3=18). For metagenomics, extracted DNA (see above) was fragmented to an average insert size of ∼350 bp using acoustic sonication (Covaris model S220). Adapter-ligated fragment libraries were generated using the Kapa Biosystems HyperPrep PCR-free library preparation workflow, according to the manufacturer’s protocol. The libraries were quantified with the KAPA qPCR library quantitation assay and sequenced on an Illumina NextSeq 500 platform with the 300 cycle Mid-Output Kit (2 × 151 bp paired end sequencing). The final output was ∼7.2 M read pairs (∼2.2 Gb) per sample.

Raw reads were filtered with BBduk. First, the last base off of 151bp reads was trimmed with “ftm=5”. Adapters were clipped off with “tbo tpe k=23 mink=11 hdist=1 ktrim=r”. PhiX sequences were filtered out with “k=31 hdist=1”. 3’ low quality bases were clipped off with “qtrim=rl trimq=15 minlength=30”. Quality-controlled reads were assembled separately for each sample and co-assembled for all samples with MEGAHIT v1.2.2-beta^47^. Contigs shorter than 500 bp were not further considered. Per contig sequencing depth was determined with BBMap v38.06 with the parameter “minid=0.99”. The coassembly and each individually assembled sample were binned separately by three methods, MetaBat v2:2.15^48^, MaxBin v2.2.7^49^ and CONCOCT v1.1.0^50^. DASTool v1.1.2 was applied to select the best bins from the three binning methods for each library^51^. dRep v3.0.0 was used to dereplicate bins obtained from different assemblies^52^. Completeness and contamination of bins (MAGs) were estimated by CheckM v1.1.3^53^. MAGs were taxonomically classified with GTDBtk v1.3.0^54^. Unbinned contigs were dereplicated by blast searches to each other. If an unbinned contig was 99% identical to a binned contig, the unbinned contig was discarded. If two unbinned contigs had 99% identity to each other, only the longer one was kept. Sequencing depth information of all non-redundant contigs were aggregated from mapping results using the script “jgi_summarize_bam_contig_depths” provided with MetaBat^48^. Contigs were annotated using MetaErg v1.2.3^55^.

The relative sequence abundance of each population associated with a MAG in metagenomes was calculated by dividing the sequencing depth of the MAG by the sum of sequencing depths of all MAGs and the unbinned contigs. Each MAG was associated with corresponding ASVs based on abundance and taxonomy. The replication rate of each population was estimated with iRep^31^. The codon usage bias of MAGs were estimated with gRodon^33^ using “Partial” mode.

### RNA extraction and metatranscriptomic sequencing

Pellets from 50 ml culture were processed for RNA extraction using the RNeasy PowerSoil Total RNA Kit (Qiagen, USA). A DNase kit (Invitrogen, CA) was used for RNA purification. The RNA concentration was checked with a Qubit 2.0 Fluorometer (Invitrogen, CA). Libraries were prepared using the New England Biolabs NEBNext rRNA depletion kit (Bacteria) and NEBNext Ultra II Directional RNA library prep kit (Illumina, San Diego, CA). The libraries were quantified by KAPA qPCR library quantitation assays and sequenced paired-end using MiSeq platform (Illumina, San Diego, CA) with a 150 cycle v3 sequencing kit, yielding ∼1.5 M reads pairs for each sample.

Read quality control was performed using the procedure described above. Reads mapping to ribosomal genes were filtered out with SortMeRNA v4.2.0^56^ with a 1×e^-10^ e-value cutoff. The filtered reads were mapped to the dereplicated contigs with 99% identity. Relative transcriptional activity for each gene was calculated based on per base sequencing depth.

To investigate transcriptional regulation of each gene in each MAG, transcriptional abundance of each gene in each MAG was normalized by total transcriptional abundance of all genes in the MAG. Relative abundance of transcripts dedicated to key metabolic processes in each MAG was calculated. Transcriptome turnover of each MAG was defined as the percentage of gene transcripts that were different between two phases or replicates. For transcriptome turnover calculations, only genes with sequencing depths of ≥ 20 in the sum of two phases or replicates were included. Transcriptome turnover for each MAG was calculated by dividing the sum of the absolute differences in normalized transcriptome sequencing depth of the genes in the MAG by two and by the number of included genes with the ‘tidyverse’ package in R v4.0.3. Transcriptome turnover values between phases and replicates were compared within and between groups in t-tests, with P-values < 0.05 considered as significant. Transcriptome turnover was also compared to expected growth of a population, assuming per population abundances did not change between phases (Supplementary Method). If the transcriptome turnover was higher than the theoretical no-change value, this was taken as evidence that a population was actively degrading old transcripts.

### Protein extraction and metaproteomics

For protein extraction, 50 mL culture pellets were transferred to lysing matrix bead tubes A (MP Biomedicals) with the addition of SDT-lysis buffer (0.1M DTT) in a 10:1 ratio^57^. Matrix tubes were bead-beated in an OMNI Bead Ruptor 24 for 45 s at 6 m s^−1^ and then incubated at 95 °C for 10 min. These steps led to pelleted, lysed cells. Peptides were isolated from pellets by filter-aided sample preparation (FASP)^58^. A Qubit 2.0 Fluorometer (Invitrogen, CA) was used to check protein concentrations. For proteomics, peptides were first separated on a 50 cm × 75 μm analytical EASY-Spray column by an an UltiMate 3000 RSLCnano Liquid Chromatograph (Thermo Fisher Scientific, Waltham, MA, USA) as previously described^59^. Eluting peptides were analyzed in a QExactive Plus hybrid quadrupole-Orbitrap mass spectrometer (Thermo Fisher Scientific, CA, USA).

Expressed proteins were identified and quantified with Proteome Discoverer version 2.0.0.802 (Thermo Fisher Scientific, CA, USA), using the Sequest HT node^57^. The Percolator Node and FidoCT were used to estimate false discovery rates (FDR) at the peptide and protein level, respectively. Peptides and proteins with FDR > 5% were discarded^59^. Relative abundance of proteins was estimated based on normalized peptide-spectral matches (PSMs). The identification database was prepared based on predicted protein sequences of all binned and unbinned contigs. Redundant proteins (> 95% amino acid identity) were removed by cd-hit^60^, while giving preference to proteins from binned contigs^57^. Phosphorylated and acetylated proteins were identified in parallel. Only unambiguous PSMs with “high” FDR confidence were included in further phosphorylation and acetylation analysis. In total, 2,216,073 MS/MS spectra were acquired, yielding 616,029 PSMs, 20,928 identified proteins and 10,655 proteins of at least “medium” confidence.

Proteomic abundance of each gene in each MAG was normalized by total protein abundance of all genes in the MAG. Relative abundance of proteins dedicated to key metabolic processes in each MAG was calculated. Proteome turnover of each MAG was defined as the percentage of proteins that were different between two phases or replicates. For proteome turnover calculations, only genes with ≥ 10 detected PSMs in the sum of two phases or replicates were included. Proteome turnover for each MAG was calculated by dividing the sum of the absolute relative protein abundance differences of the genes in the MAG by two and by the number of included genes. Proteome turnover was compared to expected growth of a population, assuming per population abundances did not change between phases (Supplementary Method). If the proteome turnover was higher than the theoretical no-change value, this was taken as evidence that a population was actively degrading old proteins. Calculation of correlation between transcriptional and translational regulation only included genes with the sum of transcript sequencing depths ≥ 60 and the sum of detected PSMs ≥ 30 in triplicates in two phases. For each MAG, the Pearson correlation coefficient between the transcriptome differences and the proteome differences of the involved genes between phases was calculated. Calculations of turnover and correlation were performed with the ‘tydiverse’ package in R v4.0.3. Turnover and correlation values were compared within and between groups in t-tests, with P-values < 0.05 considered as significant.

## Data availability

All sequences of this study, including amplicons, metagenomes, metagenome-assembled genomes and transcriptomes, are under the Bioproject PRJNA749639 (NCBI). The Biosamples of the 16S rRNA sequence are: SAMN20427124-SAMN20427234, SAMN26746688-SAMN26746699. The Biosamples for the metagenome raw reads are SAMN20395938-SAMN20395955 and the Biosamples for the metatranscriptome raw reads are SAMN20446884-SAMN20446901. The Biosamples for the MAGs are SAMN20395959-SAMN20395984. The mass spectrometry proteomics data have been deposited to the ProteomeXchange Consortium via the PRIDE partner repository^61^ with the dataset identifier PXD028583.

## Acknowledgements

The authors thank the University of Calgary’s Center for Health Genomics and Informatics for sequencing and informatics services. We thank Carmen Li for help with MiSeq sequencing, and Yihua Liu for help with R data analysis. We also thank Dan Liu and Oyeboade Adebayo for help with RNA extraction. We thank Michael Nightingale for help with sample collection. We also would like to thank Jayne Rattray, Jianwei Chen and Anirban Chakraborty for help with geochemistry analysis. This study was supported by the Natural Sciences and Engineering Research Council (NSERC) through a Discovery Grant to Marc Strous, the Canada Foundation for Innovation (CFI), the Canada First Research Excellence Fund (CFREF), the Government of Alberta, and the University of Calgary.

## Author Contributions

S.L., M.S., and M.D. designed research; S.L., D.M., A.K., and M.D. performed research; S.L., X.D., G.J., M.S., and M.D. analyzed data; S.L., M.S., and M.D. wrote the paper.

