## Supporting Information for "Frequency of change determines effectiveness of microbial response strategies in sulfidic stream microbiomes"

*Corresponding author: Muhe Diao

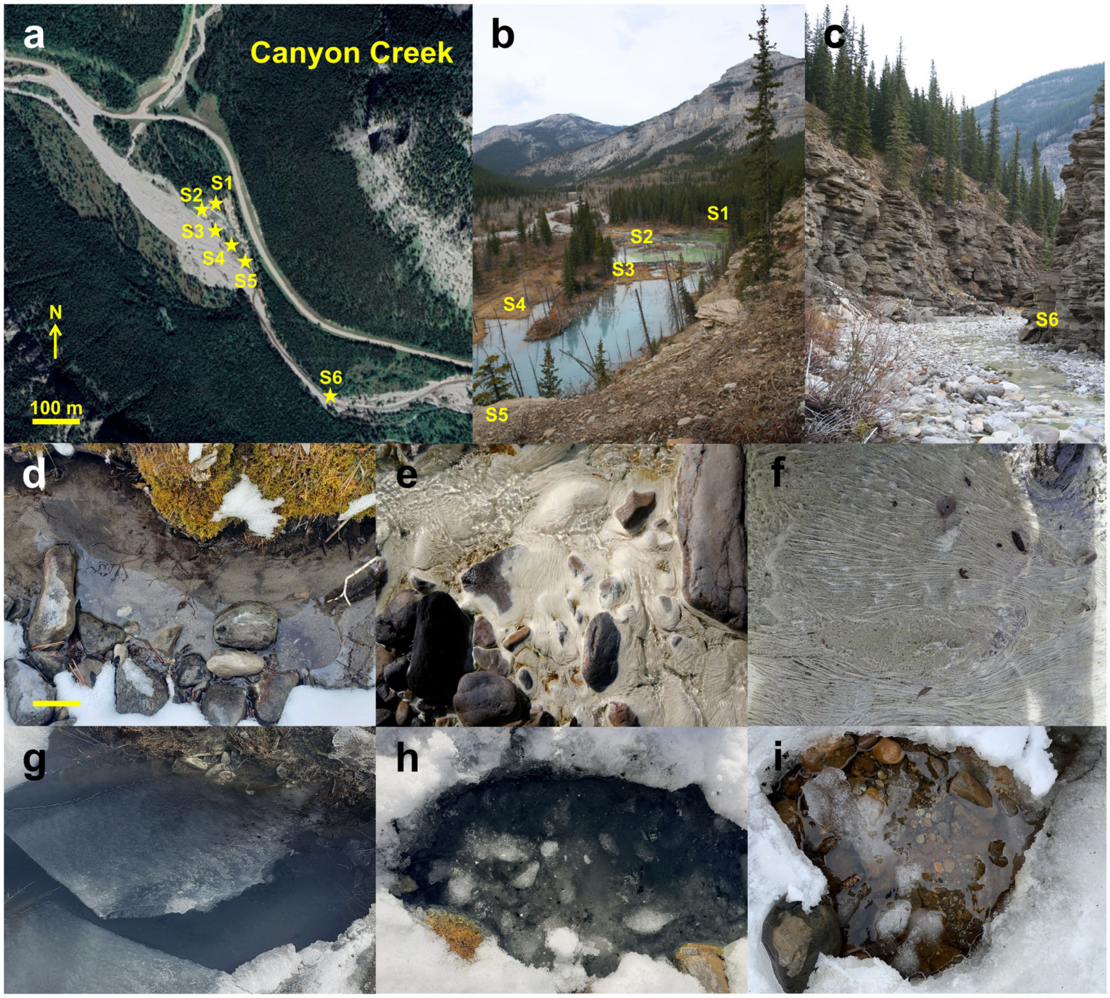


**Fig. S1 | Sampling area and sites in Canyon Creek. a**, Map of the sampling sites at Canyon Creek. The six sampling sites (S1-S6) were marked with yellow pentagrams. Map: Google Earth. **b**, Photo of the locations and surroundings of S1 to S5. **c**, Photo of the location and surroundings of S6. **d-i**, Photos of in-situ conditions of S1-S6.

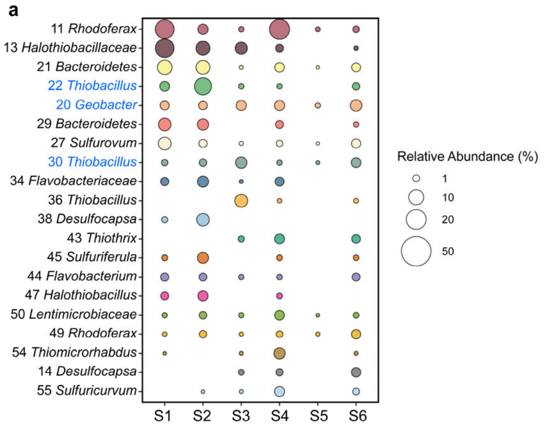


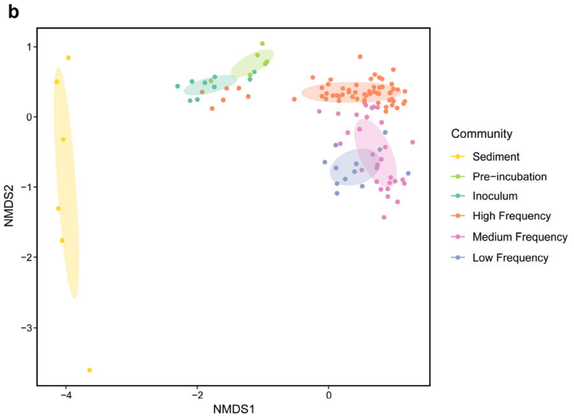


**Fig. S2 | Microbial communities of in-situ sediments and enrichment cultures. a**, Relative sequence abundances of the twenty most abundant populations (each associated with an amplicon sequence variant) in sulfidic stream sediments samples S1-S6 (Supplementary Table 2). The ASVs in blue color were represented in the chemostat incubations. **b**, NMDS (based on Bray-Curtis distances) of in-situ sediment samples, pre-incubated samples in batch culture, inoculum samples for chemostats and all samples collected along the chemostat incubations. Samples were grouped using the ‘ordiellipse’ function from the ‘vegan’ package in R.

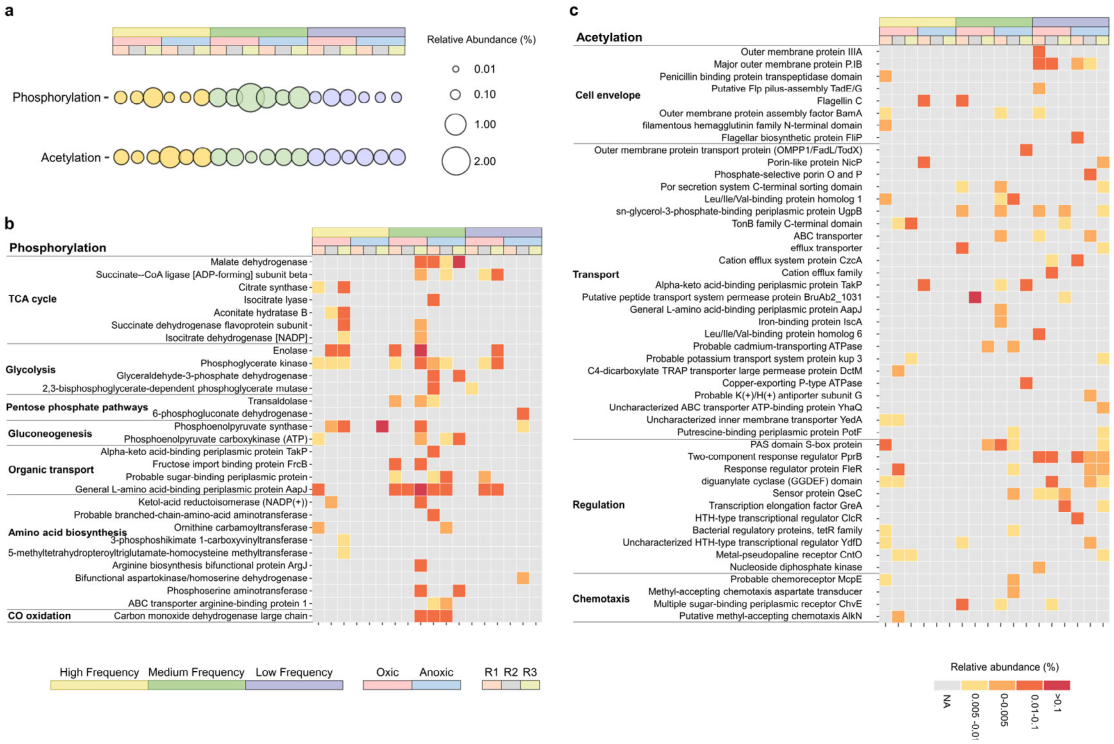


**Fig. S3 | Post-translational modification events.** **a**, Relative abundances of total phosphorylated and acetylated proteins (Supplementary Table 44). **b**, Category and relative abundances of phosphorylated proteins. **c**, Category and relative abundances of acetylated proteins (Supplementary Table 45).

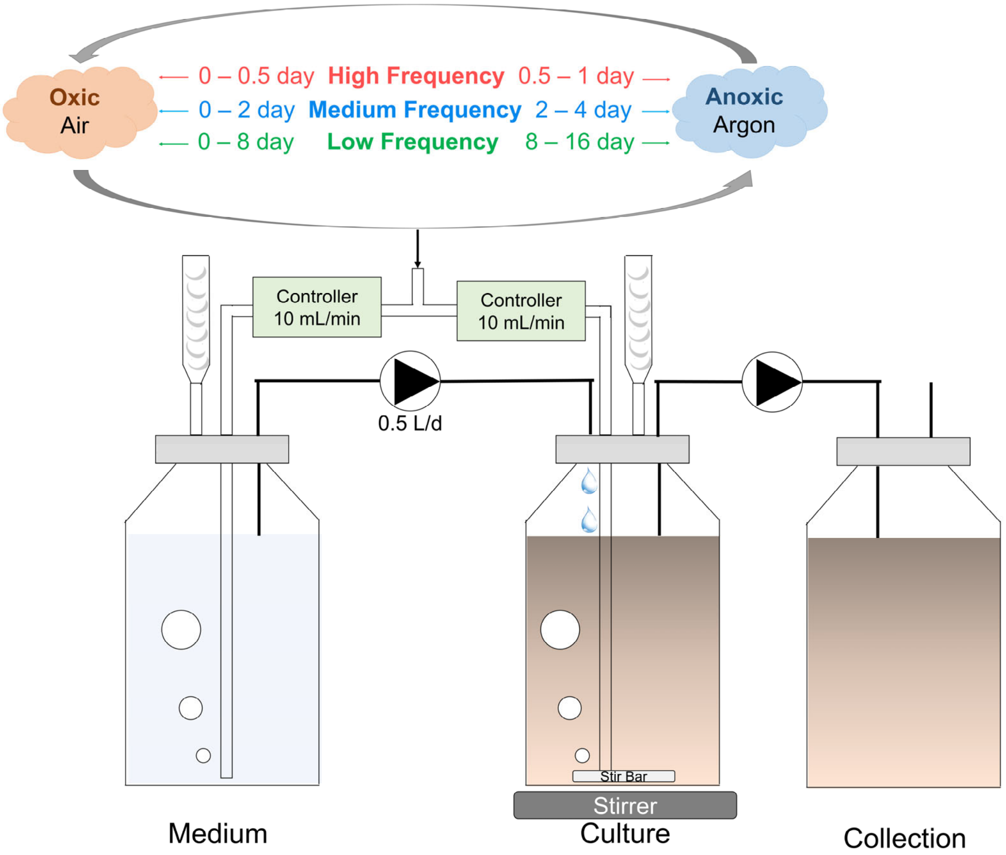


**Fig. S4 | Experimental design of chemostat cultivation.**

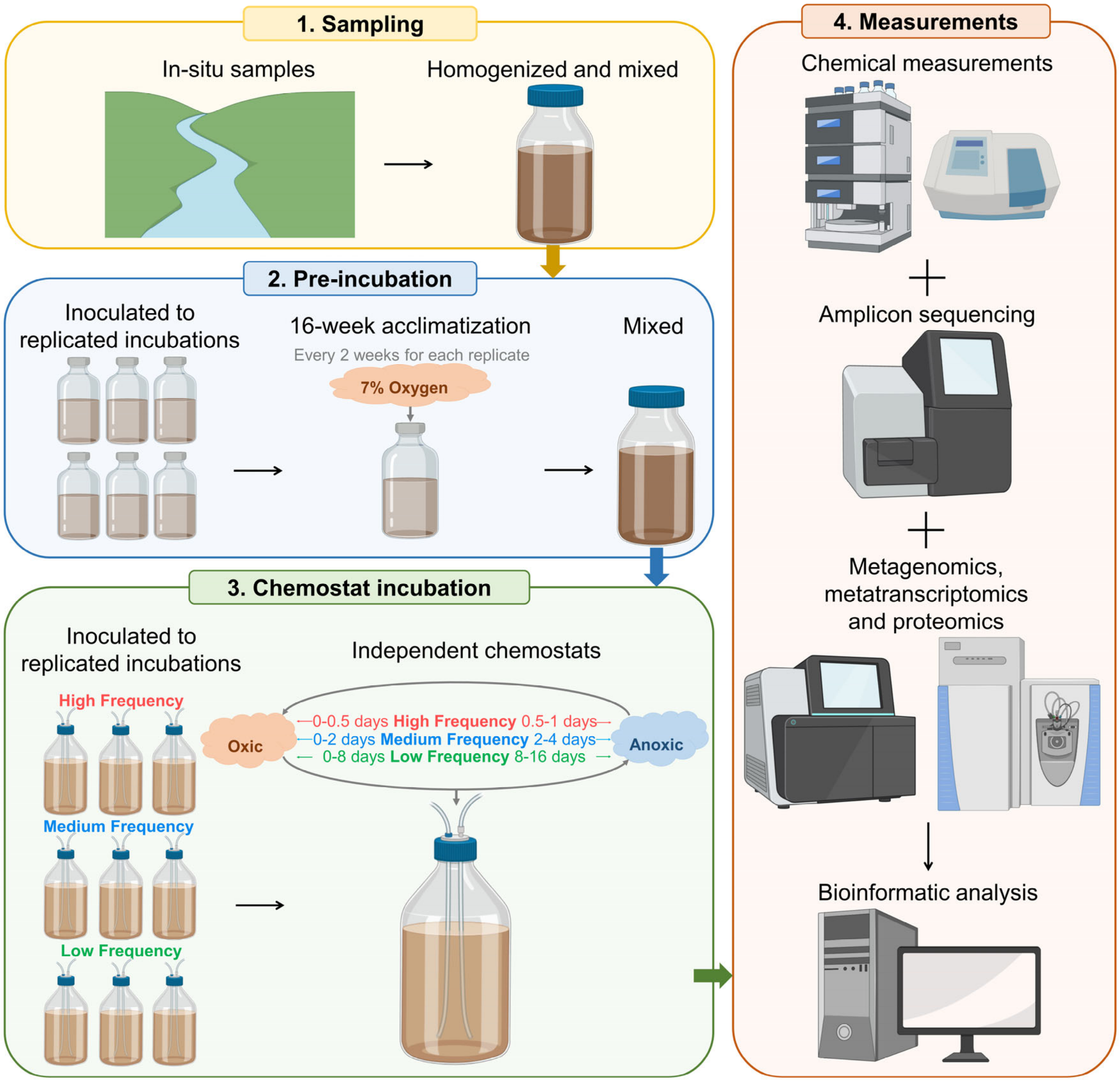


**Fig. S5 | Workflow of the experiment and data analysis.**

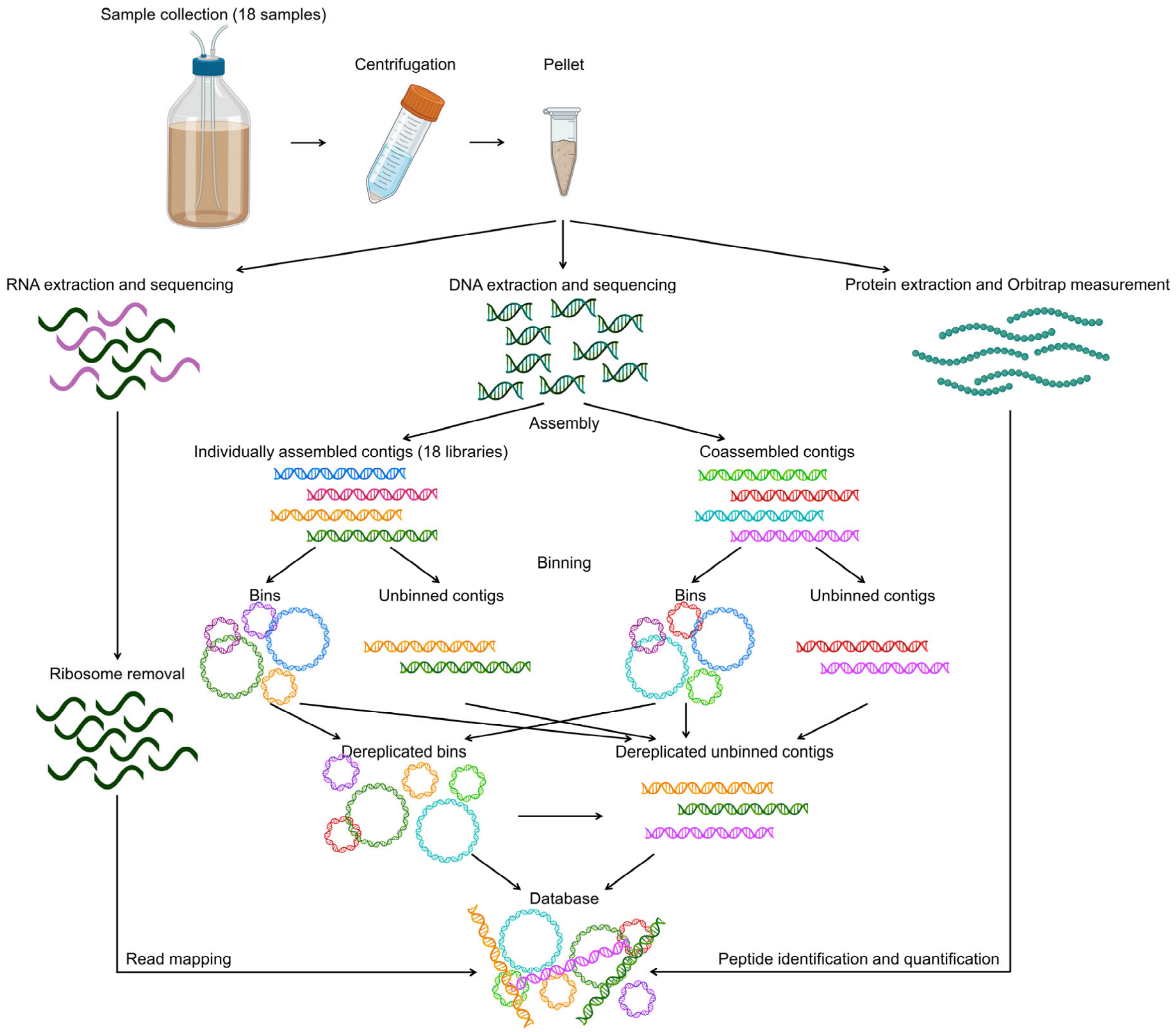


**Fig. S6 | Workflow of multi-omics analysis.**

**Supplementary Method**

In a chemostat, the mass balance of organisms is:

$$[\text{Rate of accumulation of cells}] = [\text{Rate of cells entering}] - [\text{Rate of cells leaving}] + [\text{Rate of generation of live cells}] \quad (1)$$

In our design, [Rate of cells entering] equals to zero because the medium is sterilized.

If we assume cells are not growing in the chemostat, [Rate of generation of live cells] equals to zero, and cell mass balance is worked as the followings:

$$[\text{Rate of accumulation of cells}] = V\frac{dC}{dt} \quad (2)$$

$$[\text{Rate of cells leaving}] = -vC \quad (3)$$

$$V\frac{dC}{dt} = -vC \quad (4)$$

Where $V$ is the culture volume, $C$ is concentration of cells, $v$ is the flowing rate of the effluent, $t$ is time. This can be reorganized in the following equation:

$$\frac{dC}{C} = -Ddt \quad (5)$$

Where $D$ is the dilution rate of the chemostat. In our design, per culture volume (1L) changes every 2 days, so $D$ equals to 0.5. The general solution to equation (5) is the following equation:

$$lnC = -\frac{1}{2}t + a \quad (6)$$

Where $a$ is a constant. At the beginning of the incubation, $t$ equals to zero and the cell concentration $C_o$ equals to $e^a$. At any given time, the cell concentration $C_t$ could be calculated as followings:

$$C_t = e^{-\frac{t}{2}+a} = C_o e^{-\frac{t}{2}} \quad (7)$$

$$\frac{C_t}{C_o} = e^{-\frac{t}{2}} \tag{8}$$

Assuming relative abundance of a population does not change between phases, theoretical percent of newly growing organisms of a population is calculated according to the following equation:

$$\text{Percent of newly growing organisms} = 100\% - \text{percent of remained organisms} \tag{9}$$

$$\text{Percent of newly growing organisms} = 100\% - \frac{C_t}{C_o} \tag{10}$$

$$\text{Percent of newly growing organisms} = 100\% - e^{-\frac{t}{2}} \tag{11}$$

Where $C_t$ is the concentration of cells assuming cells are not growing in the chemostat. Equation (11) is obtained based on equation (8) and (10).

At high-frequency, when $t$ is to 0.5 days, the percent of newly growing organisms of a population is 22%.

At medium-frequency, when $t$ is 2 days, the percent of newly growing organisms of a population is 63%.

At low-frequency, when $t$ is 2 days, the percent of newly growing organisms of a population is 98%.
